# Exploring the bioherbicide and biofungicide effects of pistachio hulls for their use in sustainable agriculture

**DOI:** 10.1101/2025.03.21.644490

**Authors:** A.I. González-Hernández, A. Vivar-Quintana, M.I. Saludes-Zanfaño, V.M. Gabri, M.R. Morales-Corts

## Abstract

Agriculture requires sustainable approaches for effective weed management reducing the negative consequences of synthetic herbicides. In this context, some agricultural by-products such as pistachio hulls could be considered as biopesticide compounds due to their allelopathic effect. The application of pistachio hull extract and powder produced an allelopathic effect against the weeds *Solanum nigrum, Lactuca serriola* and *Lolium rigidum* in in vitro and in vivo assays. The effect of the extract was more noticeable in the broad-leaf weeds *Solanum nigrum* and *Lactuca serriola*, while the powder seemed to be a more efficient strategy in *Lolium rigidum*. The allelopathic effects were mainly produced by the high concentration of phenolic compounds such as gallic and protocatequic acids, since the pure compounds application at the concentration found in the extract inhibited seed germination and development. Moreover, hull extract had no biofungicide effect against *Alternaria alternata, Botryosphaeria dothidea, Aspergillus niger* and *Rhizoctonia solani pathogens*. Altogether led to conclude that pistachio hull extract and powder could be a good approach to control weeds in sustainable agriculture. Further studies are required to elucidate the mode of action of these biochemicals.

## 1. Introduction

Agriculture faces up with the challenge of the need for effective weed management reducing the environmental impact of synthetic herbicides. The use of conventional herbicides had led to adverse effects on biodiversity, soil health, and water quality [1]. Furthermore, the emergence of herbicide-resistant weed species has further complicated weed management strategies, prompting a search for sustainable alternatives [2]. The latest European regulations address a reduction of 50% in the application of synthetic pesticides by 2030, making natural herbicides or bioherbicides a promising alternative. These products are derived from natural sources, such as plants, microorganisms, or minerals, and are often perceived as safer for the environment and human health [3, 4). In this context, some agricultural by-products have been shown to have an allelopathic effect that makes them potential candidates for a role as bioherbicides. Plant by-products could be included in crop rotations or directly applied as a sustainable herbicide [5]. Their application for this purpose would allow a more sustainable management of weed control while reducing waste production [6].

Allelopathy is defined as a biological and naturally occurring phenomenon in which one plant has direct or indirect effects on another plant by the release of chemical compounds (allelochemicals) into the environment [7]. The mentioned allelochemicals are mainly alcohols, fatty acids, phenolic compounds, flavonoids, terpenoids and steroids, with the phenolic compounds being the most identified. Phenolic compounds are one of the main secondary metabolites playing a role in physiology and morphology of plants, being flavonoids and anthocyanins, a group of polyphenolic substances with a wide range of biological functions as protection against plant stress [8]. The allelopathic potential of different plant species has been studied [9] and effects ranging from inhibitory to stimulating depending on the concentration and environmental context has been found [10].

Pistachio hulls constitute between 35% and 45% of the waste generated by the pistachio producers, which can lead to environmental problems [11]. In this sense, the inhibitor effect of different parts of the pistachio tree on weeds has been previously reported. Alyousef and Ibrahim [12] showed that pistachio hulls and leaves inhibited the growth of *Diplotaxis erucoides, Sonchus arvensis* and *Papaver hybridum*. Likewise, *Pistacia khinjuk, Pistacia vera* and *Pistacia atlantica* spp. leaf extracts also had an allelopathic effect on the germination of a wide range of weeds such as *Amaranthus retroflexus, Lolium rigidum* and *Lactuca serriola* [13-15]. In the case of pistachio, compounds with allelopathic effect have been described in roots, leaves and shells of pistachio cultivars, being their concentration higher in shells and leaves [15-17]. The most predominant compounds were gallic acid, vanillic acid, p-coumaric acid, naringenin, or catechin, among others [16, 18-20]. The hulls are a great source of bioactive compounds such as antioxidants and phenolic compounds, which could play a pivotal role in weed management by their allelopathic effect [21]. Nevertheless, more studies are required to elucidate the effect of pistachio hulls as bioherbicides against the most common weeds.

Allelopathy also includes antifungal activity, since the chemical effects that a plant exerts on organisms in its environment include not only other plants, but also microorganisms such as fungi and bacteria. Among the biotic stresses, phytopathogenic fungi reduce crop yield and quality and cause important annual losses in crops by producing diseases in commercial crops and causing certain problems in the food supply chain [22]. As mentioned above, plants are capable of synthesizing secondary metabolites, which can be extracted using techniques conventional and non-conventional for the formulation of biofungicides [23]. Thus, plants such as eucalyptus and species of the Brassicaceae family release compounds such as essential oils or glucosinolates that have demonstrated antifungal activity [24]. Several studies have highlighted the antifungal efficacy of extracts from pistachio hulls, which can inhibit the growth of pathogenic fungi such as *Aspergillus flavus* [25]. The use of these natural fungicides not only minimizes chemical residues in the environment but also adds value to agricultural waste, contributing to circular economy principles.

The present study delves into the potential of pistachio hulls as natural herbicide in *Lactuca serriola, Solanum nigrum* and *Lolium rigidum* weeds, and as fungicide against *Alternaria alternata, Botryosphaeria dothidea, Aspergillus niger* and *Rhizoctonia solani* fungi, exploring their chemical composition and phytotoxic effects.

## 2. Materials & Methods

### 2.1. Preparation of the solid and liquid hull-based bioherbicide

Pistachio hulls of five-years old *Pistacia vera* L. (cv. Kerman) grown in an orchard located in the village of Parada de Rubiales (Salamanca-Castilla y León-Spain) (41_9018.0200 N, 5_26050.2400 W, 844 m.a.s.l.) were collected. Hulls were dried at 40 ºC for 9 hours in a stove (Indelab, Spain) and powdered with a grinder (Moulinex, Spain). The powder was divided in two sets: the first one that was stored at -20ºC until use and the second one that was used for extract preparation. The extract was prepared by mixing the powder with distilled water in a proportion 10:90 (w/w). This mix was soaked in darkness for 24 h and then filtered with filter paper. Once the extract was obtained, a dilution was carried out to get a more diluted extract (2.5%). Both extracts were kept at 5 ºC in the fridge until use.

### 2.2. In vitro assays in weeds and crops

The *in vitro* bioassays were performed in Petri dishes to analyse the possible inhibition effect of both hull extracts in the germination of different common weeds of the Mediterranean region: *Lactuca serriola* (broad-leaf weed), *Lolium rigidum* Gaud. (narrow-leaf weed), and *Solanum nigrum* (broad-leaf weed). Furthermore, the study of the effect on four staple crops were analysed in order to check that there is no bioherbicidal effect on them. The studied crops were: *Zea mays* L. (corn) cv. Celestquatro, *Hordeum vulgare* L. (barley) cv. Yurico, *Triticum aestivum* L. (wheat) cv. Rimbaud, and *Lens culinaris* L. (lentil) cv. Castellana. Fifteen seeds of each species were placed in a 90-mm Petri Dish with a double filter paper on the bottom soaked with 10 mL of the corresponding treatment. The considered treatments were: 1) Control treatment (distilled water), 2) Hull extract 2.5%, and 3) Hull extract 10%. Petri dishes containing seeds were placed in a growth chamber at 24/10 ºC (day/night), 16/8 h (day/night) for 8 days, except Solanum nigrum Petri dishes that were placed at 28/18 ºC (day/night), 12/12 h (day/night) for 10 days. Once the seeds germinated, the germination percentage, germination index, and radicle and epicotyl lengths were determined. The germination percentage was calculated by dividing the germinated seeds by the total seed number, multiplying by 100. The germination index (GI) was determined by dividing the corresponding treatment germination rate by the control germination rate. Four replicates of each condition were performed.

### 2.3. Pot assays

To further analyse the effect of the extract 10% and the ground hull as a preemergence herbicide in maize and lentil crops, a pot experiment was performed. Vermiculite containing pots were prepared and fertilized with a dose of the fertilizer Fertilent 20:12:8 of 300 kg ha^-1^ for maize pots and 150 kg ha^-1^ for lentil pots, in order to avoid nutrient deficiency. Then, two seeds of the crop (maize or lentil) and fifteen seeds of each of the three studied weed species were placed in each pot (2 l). Once the seeds were established in the substrate, three different treatments were considered: control (application of 10 mL of distilled water), extract 10% (application of 10 mL of hull extract 10%) and ground hull (application of 1 g of powdered hulls). Treatments were applied two times, at sowing and fifteen days later. The liquid treatments were distributed using a Cofan sprayer. Plants were irrigated when needed, applying the same volume of dechlorinated water to all the pots via drenching to avoid bioherbicide lixiviation. At the middle of the cycle, a new dosage of fertilizer was provided. Forty-five days later, the following parameters of each crop and weed species were recorded: fresh weight, dry weight and plant number. In lentil and maize assays, plants were grown under the following greenhouse conditions: temperature of 24/18 ºC (day/night) and 12/12 h photoperiod. Six replicates of each assay were performed.

### 2.4. Determination of the phenolic compounds

The analysis and identification of the polyphenols, of the liquid hull-based bioherbicide, was carried out by HPLC-Q-TOF (Agilent 6520 Accurate-Mass Q-TOF LC/MS System with Agilent 1200 Series HPLC, Arcade, USA). The separation of the compounds was carried out using a Zorbax Eclipse Plus C18 column (100 mm x 4.6 mm, 3.5 μm Agilent, USA and using as mobile phase a 0.1 % aqueous solution of formic acid (Eluent A) and acetonitrile with 0.1 % formic acid (Eluent B). Elution is carried out at a flow rate of 0.350 mL/min, under gradient conditions as follow: 0–10 min, 10% B; 10–40 min, 20% B; 40–60 min, 60 % B; 60–80 min, 90% B; 80–85 min, 90% B. Compounds have been identified by comparing the theoretical exact mass with the measured exact mass and with a maximum error tolerance in ppm equal to or less than 10 ppm. Data was acquired in the negative mode from *m/z* 150 to 1000. The gallic acid and protocatechuic acid content of the liquid hull-based bioherbicide was quantified with standard patterns of both compounds (Merck, Spain). For this purpose, a standard curve for each of the compounds was obtained in a range between 2.5 mg/l and 30 mg/l.

### 2.5. Germination bioassays with the pure phenolic compounds

The concentration of the phenolic compounds obtained in the 10% hull extract was used to prepare the pure phenolic solutions. Gallic acid and protocatechuic acid were the most predominant compounds in the extract. Thus, three different treatments were tested: control (distilled water), gallic acid (2.38 mg ml^-1^) and protocatechuic acid (4.14 mg ml^-1^). 60-mm Petri dishes with a filter paper at the bottom were used to carry out this assay. Fifteen seeds of the corresponding weed species (*Lactuca serriola, Lolium rigidum* or *Solanum nigrum*) were placed on the top of the filter paper and 1 ml of the treatment was applied dampening the filter paper. Petri dishes containing seeds were placed in a growth chamber at 24/10 ºC (day/night), 16/8 h (day/night) for 8 days, except Solanum nigrum Petri dishes that were placed at 28/18 ºC (day/night), 12/12 h (day/night) for 10 days. Once the seeds germinated, the germination percentage, germination index, and radicle and epicotyl lengths were determined as indicated in the section 2.2. Three replicates were performed.

### 2.6. Biofungicide effect of the hull extract

To study the biofungicide effect of pistachio hull an extract was prepared by mixing the powder with distilled water in a proportion 25:75 (w/w). This mix was also soaked in darkness for 24 h and then filtered with filter paper. An *in vitro* assay with *Alternaria alternata, Aspergillus niger* and *Botryosphaeria dothidea* (strains 13E062, 12L082 and 14D100 from the culture collection of UC Davis, California, USA) and *Rhizoctonia solani* (strain 122 from the collection of the Regional Center for Pest and Diseases Diagnosis, Junta de Castilla y León, Spain) was performed in sterilized potato dextrose agar (PDA) medium. The PDA medium was prepared dissolving 19.5 grams in 500 ml of water. Two treatments were considered: 1) control with sterilized distilled water, and 2) 25% filtered hull extract. Fifty µl of the corresponding treatment was added to a well located at 1 cm of the outer of the 100 × 15 mm Petri dishes (VWR Scientific Petri Dishes). Extracts were firstly sterilized by filtration through a 0.22 µm nylon filter. Then, a plug of 5 mm of diameter of the corresponding fungi was placed in the middle of the medium. The Petri dishes were kept at 27ºC for 6 days. After this time, the mean area of each pathogen was assessed determining the growth area by using ImageJ software. Five replicates of each treatment and pathogen were performed.

### 2.7. Statistical analyses

Statistical analyses were carried out with Statgraphics Centurion XVIII software (Statistical Graphics Corp., Rockville, MD, USA). The results included means with standard errors, and Tukey’s honest significant difference (HSD) post hoc test with a 95% confidence interval (p < 0.05) was performed to compare the individual means of the three considered treatments when ANOVA test found significant differences. In case of only two treatments t-Student test was carried out.

## 3. Results

### 3.1. *In vitro* assay in weeds and crops

The effect of both hull’s concentration extracts was tested in vitro to study the bioherbicide effect at germination stage against the weeds *Lactuca serriola, Solanum nigrum* and *Lolium rigidum* (Table 1) and the crops *Zea mays, Hordeum vulgare, Triticum aestivum* and *Lens culinaris* (Table 2). The results showed a significant reduction in germination and radicle and epicothyl growth of *Lactuca serriola* under the 2.5% and 10% extract treatments compared to the control treatment. No growth or germination was observed when 10% extract was applied. In the case of *Solanum nigrum*, a total reduction in germination and radicle and epicotyl length was observed under both extract concentrations. In addition, the narrow-leaf species *Lolium rigidum* showed a reduction in the epicotyl and radicle length after the application of both extracts, being more noticeable after the application of the 10% extract. However, the germination of this species was not affected by the 2.5% extract.

**Table 1.** *In vitro* effect of hull’s extracts on the three weeds germination.

| Weeds | Treatment | Epicotyl length (mm) | Radicle length (mm) | Germination percentage | GI |
| --- | --- | --- | --- | --- | --- |
| *Lactuca serriola* | Control | 10.40±1.68 a | 7.64±1.29 a | 55.56±7.59 a | 1 |
|  | Extract 2.5% | 3.16±1.17 b | 1.13±0.62 b | 17.78±4.59 b | 0.32 |
|  | Extract 10% | 0.00±0.00 b | 0.00±0.00 b | 0.00±0.00 c | 0 |
| *Solanum nigrum* | Control | 0.69±0.04 a | 1.81±0.14 a | 93.33±0.00 a | 1 |
|  | Extract 2.5% | 0.00±0.00 b | 0.00±0.00 b | 0.00±0.00 b | 0 |
|  | Extract 10% | 0.00±0.00 b | 0.00±0.00 b | 0.00±0.00 b | 0 |
| *Lolium rigidum* | Control | 49.66±3.15 a | 59.96±4.30 a | 88.89±1.52 a | 1 |
|  | Extract 2.5% | 39.30±3.84 b | 15.67±1.88 b | 88.89±2.07 a | 1 |
|  | Extract 10% | 5.30±1.69 c | 0.74±0.21 c | 28.89±2.50 b | 0.33 |

Different letters in the same column mean statistically significant differences $p < 0.05$ , according to Tukey's HSD test

**Table 2.** *In vitro* effect of hull’s extracts on the four crops germination.

| Crop | Treatment | Epicotyl length (mm) | Radicle length (mm) | Germination percentage | GI |
| --- | --- | --- | --- | --- | --- |
| *Zea mays* | Control | 39.40±1.68 a | 53.62±2.55 a | 97.78±0.57 a | 1 |
|  | Extract 2.5% | 28.18±4.22 b | 35.67±3.29 b | 88.89±1.15 a | 0.91 |
|  | Extract 10% | 18.84±1.86 c | 13.51±1.50 c | 75.56±1.15 b | 0.77 |
| *Hordeum vulgare* | Control | 64.42 ± 7.03 a | 50.64±2.41 a | 97.78±0.57 a | 1 |
|  | Extract 2.5% | 55.29±4.28 a | 26.09±2.08 b | 95.55±0.57 a | 0.98 |
|  | Extract 10% | 1.22±0.67 b | 1.31±0.67 c | 82.22±4.59 a | 0.84 |
| *Triticum aestivum* | Control | 54.42±2.76 a | 50.24±2.99 a | 97.78±0.57 a | 1 |
|  | Extract 2.5% | 33.64±3.49 b | 17.67±2.19 b | 82.22±2.30 a | 0.84 |
|  | Extract 10% | 2.96±0.99 c | 1.27±0.61 c | 20.00±0.99 b | 0.20 |
| *Lens culinaris* | Control | 33.10±2.78 a | 22.47±1.74 a | 88.89±2.87 a | 1 |
|  | Extract 2.5% | 22.86±2.03 b | 17.06±1.44 b | 88.89±2.87 a | 1 |
|  | Extract 10% | 2.34±0.72 c | 1.07±0.32 c | 26.67±6.89 b | 0.30 |

Different letters in the same column mean statistically significant differences $p < 0.05$ , according to Tukey's HSD test

Concerning crops, all species showed a significant reduction in epicotyl and radicle length under the two extract concentrations compared to the control treatment, except for the epicotyl length in *Hordeum vulgare*, showing similar results under 2.5% extract and control treatments. Regarding the germination percentage, the 2.5% extract did not significantly reduce this parameter in any of the crop species, while the 10% extract significantly reduced the germination percentage in corn, wheat and lentil.

### 3.2. *In vivo* assay

To further study the effect of the extract and the solid and ground hulls, a pot assay with two different crops was carried out. The effect of hull extract and the dry powder (ground hulls) on lentil (Table 3) and corn (Table 4) with a total of 30 seeds of the three weeds was analyzed. The results showed that the application of both hull-based bioherbicides reduced by around 90% the number of germinated plants, the fresh weight and the dry weight of *Solanum nigrum* in both the lentil assay (Table 3) and the corn assay (Table 4). In *Lactuca serriola*, both bioherbicide formats reduced the parameters studied, although under the dry powder application no significant differences in the number of plants and the fresh weight in the lentil test were observed (Table 3). Curiously, when the growth of this weed was studied in the corn assay, no significant differences were observed between treatments (Table 4), but the values of the control were lower than those observed in the lentil crop. Finally, for the narrow-leaf species *Lolium rigidum*, a reduction in fresh and dry weight was observed in both assays with the application of the bioherbicide in powder form despite no differences were found in the number of plants. It should be noted that no significant differences were found in crops (*Lens culinaris* and *Zea mays*).

**Table 3.** Bioherbicide effect on lentil crop and weeds.

| Plant species | Treatment | Fresh weight (g) | Dry weight (g) | Plant number |
| --- | --- | --- | --- | --- |
| *Lens culinaris* | Control | 2.77±0.21 b | 0.45±0.03 a |  |
|  | Extract | 2.82±0.19 b | 0.42±0.03 a |  |
|  | Ground hull | 3.66±0.21 a | 0.47±0.02 a |  |
| *Solanum nigrum* | Control | 0.41±0.08 a | 0.06±0.01 a | 8.33±0.88 a |
|  | Extract | 0.05±0.04 b | 0.01±0.00 b | 0.67±0.21 b |
|  | Ground hull | 0.07±0.06 b | 0.01±0.01 b | 0.83±0.65 b |
| *Lactuca serriola* | Control | 2.06±0.15 a | 0.21±0.02 a | 6.00±0.52 a |
|  | Extract | 0.24±0.05 b | 0.03±0.00 b | 2.50±0.43 b |
|  | Ground hull | 1.29±0.57 a | 0.10±0.04 b | 3.83±1.14 ab |
| *Lolium rigidum* | Control | 19.85±1.66 a | 2.17±0.19 a | 12.17±0.79 a |
|  | Extract | 17.55±1.27 ab | 1.82±0.14 ab | 12.50±0.62 a |
|  | Ground hull | 15.24±1.22 b | 1.50±0.13 b | 11.50±0.56 a |

Different letters in the same column mean statistically significant differences $p<0.05$ , according to Tukey's HSD test

**Table 4.** Bioherbicide effect on corn crop and weeds.

| Plant species | Treatment | Fresh weight (g) | Dry weight (g) | Plant number |
| --- | --- | --- | --- | --- |
| *Zea mays* | Control | 20.20±1.34 a | 2.04±0.15 a |  |
|  | Extract | 19.45±0.82 a | 1.87±0.09 a |  |
|  | Ground hull | 17.15±1.60 a | 1.86±0.16 a |  |
| *Solanum nigrum* | Control | 0.38±0.09 a | 0.06±0.02 a | 5.83±0.87 a |
|  | Extract | 0.00±0.00 b | 0.00±0.00 b | 0.17±0.17 b |
|  | Ground hull | 0.08±0.03 b | 0.01±0.00 b | 1.50±0.56 b |
| *Lactuca serriola* | Control | 0.51±0.29 a | 0.06±0.04 a | 2.33±1.28 a |
|  | Extract | 0.48±0.22 a | 0.06±0.03 a | 2.33±0.33 a |
|  | Ground hull | 0.47±0.18 a | 0.05±0.03 a | 2.33±0.71 a |
| *Lolium rigidum* | Control | 15.69±1.00 a | 1.69±0.12 a | 11.50±0.76 a |
|  | Extract | 14.46±1.16 a | 1.51±0.14 a | 11.17±0.40 ab |
|  | Ground hull | 10.57±0.77 b | 1.13±0.08 b | 9.67±0.56 b |

Different letters in the same column mean statistically significant differences $p<0.05$ , according to Tukey's HSD test

### 3.3. Determination of the phenolic compounds in pistachio hulls

The 10% aqueous extract was analyzed for its phenolic composition. A total of 18 compounds belonging to eight different families were identified (Table 5). Within the family of hydroxybenzoic acids 5 compounds were identified (Theogallin, Digallic acid, Gallic acid, protocatechuic acid, Vanillic acid hexoside and Ellagic acid), two compounds in the family of hydroxynamic acids (p-Coumaric acid and o-Coumaric acid) and two tannins belonged to the family of galloyl derivatives (Mono-galloyl-glucoseI and Galloyl dihexose). Three flavonones (eriodictyol hexoside, eriodictyol and naringenin), two flavones (luteolin and apigenin), one flavanonols (taxifolin) and one flavan-3-ol were also identified. Finally, one anthocyanin, cyanidin 3-O-glucoside, responsible for the reddish color of the skins, was identified.

**Table 5.**
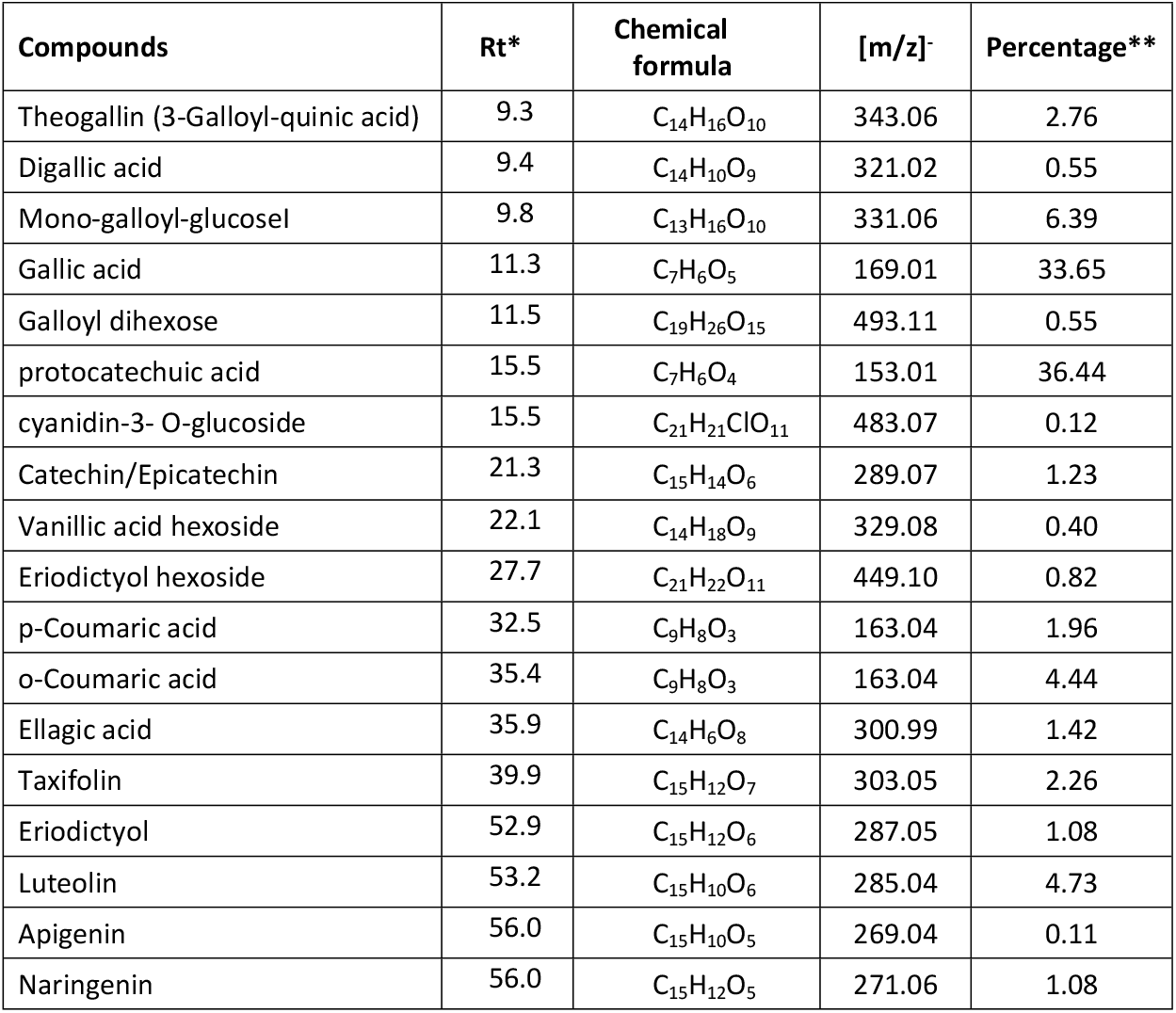
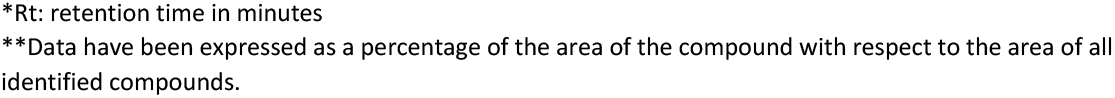
Chemical profile of the identified phenolic compounds in pistachio hull by HPLC-Q-TOF.

| Compounds | Rt* | Chemical formula | [m/z] | Percentage** |
| --- | --- | --- | --- | --- |
| Theogallin (3-Galloyl-quinic acid) | 9.3 | C<sub>14</sub>H<sub>16</sub>O<sub>10</sub> | 343.06 | 2.76 |
| Digallic acid | 9.4 | C<sub>14</sub>H<sub>10</sub>O<sub>9</sub> | 321.02 | 0.55 |
| Mono-galloyl-glucosel | 9.8 | C<sub>13</sub>H<sub>16</sub>O<sub>10</sub> | 331.06 | 6.39 |
| Gallic acid | 11.3 | C<sub>7</sub>H<sub>6</sub>O<sub>5</sub> | 169.01 | 33.65 |
| Galloyl dihexose | 11.5 | C<sub>19</sub>H<sub>26</sub>O<sub>15</sub> | 493.11 | 0.55 |
| protocatechuic acid | 15.5 | C<sub>7</sub>H<sub>6</sub>O<sub>4</sub> | 153.01 | 36.44 |
| cyanidin-3- O-glucoside | 15.5 | C<sub>21</sub>H<sub>21</sub>ClO<sub>11</sub> | 483.07 | 0.12 |
| Catechin/Epicatechin | 21.3 | C<sub>15</sub>H<sub>14</sub>O<sub>6</sub> | 289.07 | 1.23 |
| Vanillic acid hexoside | 22.1 | C<sub>14</sub>H<sub>18</sub>O<sub>9</sub> | 329.08 | 0.40 |
| Eriodictyol hexoside | 27.7 | C<sub>21</sub>H<sub>22</sub>O<sub>11</sub> | 449.10 | 0.82 |
| p-Coumaric acid | 32.5 | C<sub>9</sub>H<sub>8</sub>O<sub>3</sub> | 163.04 | 1.96 |
| o-Coumaric acid | 35.4 | C<sub>9</sub>H<sub>8</sub>O<sub>3</sub> | 163.04 | 4.44 |
| Ellagic acid | 35.9 | C<sub>14</sub>H<sub>6</sub>O<sub>8</sub> | 300.99 | 1.42 |
| Taxifolin | 39.9 | C<sub>15</sub>H<sub>12</sub>O<sub>7</sub> | 303.05 | 2.26 |
| Eriodictyol | 52.9 | C<sub>15</sub>H<sub>12</sub>O<sub>6</sub> | 287.05 | 1.08 |
| Luteolin | 53.2 | C<sub>15</sub>H<sub>10</sub>O<sub>6</sub> | 285.04 | 4.73 |
| Apigenin | 56.0 | C<sub>15</sub>H<sub>10</sub>O<sub>5</sub> | 269.04 | 0.11 |
| Naringenin | 56.0 | C<sub>15</sub>H<sub>12</sub>O<sub>5</sub> | 271.06 | 1.08 |

\*Rt: retention time in minutes
\*\*Data have been expressed as a percentage of the area of the compound with respect to the area of all identified compounds.

The contribution of each of these compounds in the extracts was established from their area in the chromatograms. Thus, it was observed that hydroxybenzoic acids represent more than 75% of the total area, being the most abundant group mainly due to gallic acid and protocatechuic acids. Given the importance of both compounds, a quantification of their concentration in the extracts was carried out using commercial standards as indicated in section 2.4. Thus, it was obtained that the concentration presented in the 10% pistachio hull extract, amounted to 13.33 g/l of gallic acid (± 4.73) and 4.13 g/l of protocatechuic acid (± 2.92).

### 3.4. Pure phenolic compounds effect in seed germination and development

Gallic and protocatechuic acids were tested to evaluate their individual effects by biogermination assays. The amounts assayed were those quantified in the 10% extracts. The results showed that the germination process of the studied weeds has been reduced by both gallic acid and protocatechuic acid (Table 6). These phenolic compounds produced a significant inhibition of the epicotyl and radicle length in the studied narrow and broad leaf weeds. Nevertheless, the exposure of *Lolium rigidum* to protocatechuic acid reduced epicotyl length compared to gallic acid. No significant differences were observed in the germination percentage.

**Table 6.** Pure phenolic compounds effect on seed germination and development.

| Weeds | Treatment | Epicotyl length (cm) | Radicle length (cm) | Germination percentage | GI |
| --- | --- | --- | --- | --- | --- |
| *Lolium rigidum* | Control | 2.72 ± 0.26 a | 2.72 ± 0.28 a | 81.4 ± 9.69 a | - |
|  | Gallic acid | 1.64 ± 0.31 b | 0.07 ± 0.03 b | 53.33 ± 23.33 a | 0.66 |
|  | Protocatequic acid | 0.38 ± 0.16 c | 0.00 ± 0.00 b | 23.00 ± 14.14 a | 0.28 |
| *Lactuca serriola* | Control | 0.68 ± 0.13 a | 0.66 ± 0.11 a | 58.67 ± 16.25 a | - |
|  | Gallic acid | 0.07 ± 0.02 b | 0.24 ± 0.05 b | 50.00 ± 10.00 a | 0.85 |
|  | Protocatequic acid | 0.00 ± 0.00 b | 0.08 ± 0.02 b | 53.33 ± 0.00 a | 0.91 |
| *Solanum nigrum* | Control | 0.87 ± 0.13 a | 0.93 ± 0.19 a | 52.00 ± 16.79 a | - |
|  | Gallic acid | 0.07 ± 0.02 b | 0.24 ± 0.05 b | 20.00 ± 10.00 a | 0.96 |
|  | Protocatequic acid | 0.00 ± 0.00 b | 0.14 ± 0.01 b | 90.00 ± 4.71 a | 1.73 |

Different letters in the same column mean statistically significant differences $p < 0.05$ , according to Tukey's HSD test

### 3.5. Biofungicide effect of pistachio hull extract against *Alternaria alternata, Botryosphaeria dothidea, Aspergillus niger* and *Rhizoctonia solani*

To further understand the effect of the hull extract an *in vitro* experiment was carried out to determine the biofungicide effect against *Alternaria alternata, Botryosphaeria dothidea, Aspergillus niger* and *Rhizoctonia solani* (Figure 1). No effect in the reduction of the pathogen’s growth was observed in the petri dishes except a slight decrease in *Botryosphaeria dothidea* (Figure 1). No significant differences were observed in the *in vitro* fungi growth when hull extracts were applied.

**Figure 1.**
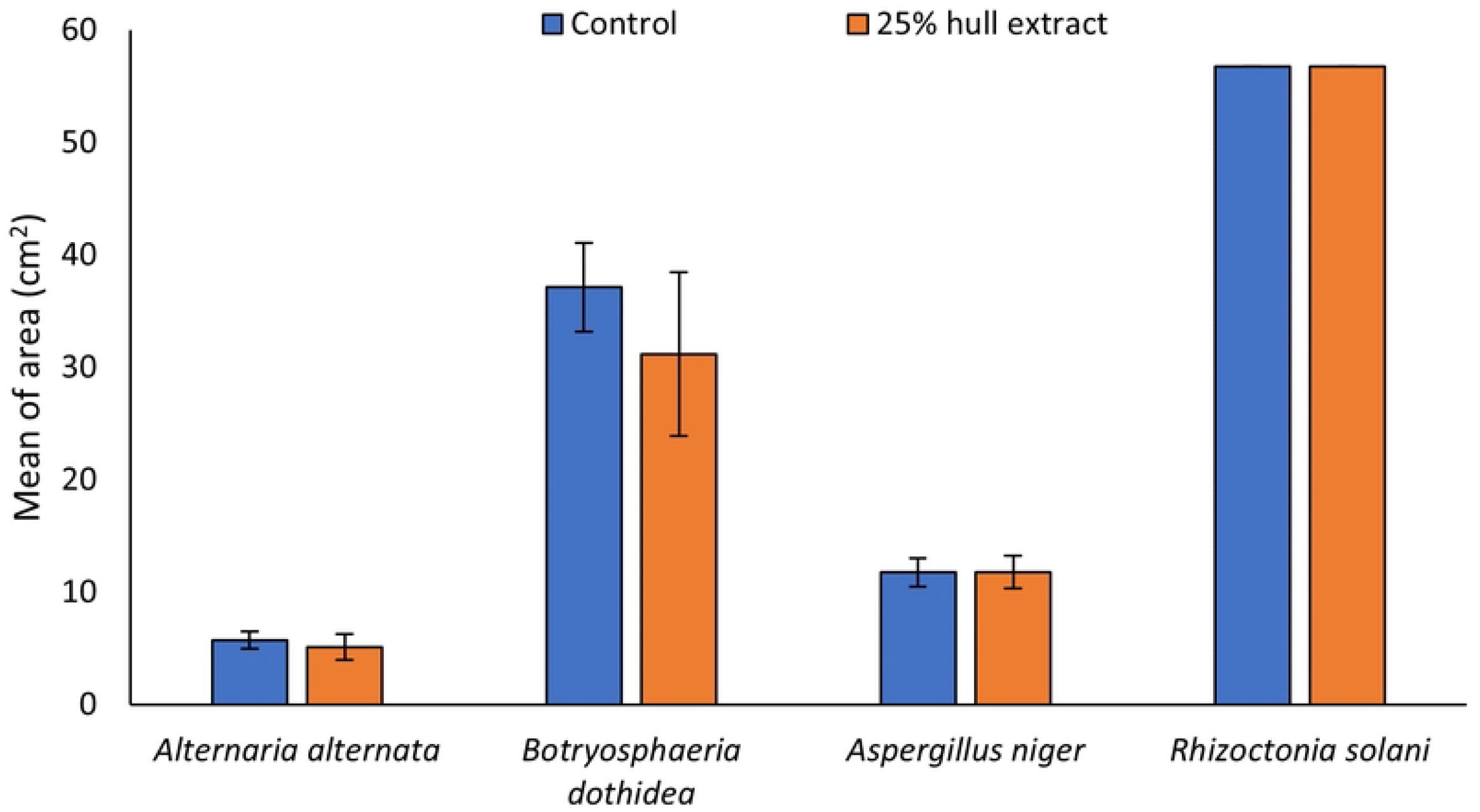
Biofungicide effect of the 25% hull extract against *Alternaria alternata. Botryosphaeria dothidea. Aspergillus niger* and *Rhizoctonia solani*.

## 4. Discussion

The agricultural sector produces a large volume of waste that is unused productively, leading to environmental problems, economic costs and loss of valuable compounds, among others [26]. This waste can be valorised for reuse in crop production because of its high levels of bioactive compounds [27]. These natural ingredients are increasingly being seen as potential solutions to combat weed resistance and address the environmental issues associated with synthetic herbicides [28]. Bioherbicides are products naturally originated which can be used to control weeds that compete with crops by light, water and nutrientes [29]. Those bioherbicides derived from allelochemicals are eco-friendly, as they have a short environmental lifespan, minimizing the risk of soil and water contamination while posing no significant harm to non-targeted organisms [29, 30]. Thus, the use of bioherbicides constitutes a proactive step towards a more sustainable and environmentally friendly future for European agriculture offering a viable strategy to decrease reliance on chemical fungicides and pesticides. In this study, we focus on the interest on the bioherbicidal effects of *Pistacia vera* hull, extract and powder, on three common weed species in the Mediterranean crop fields, as well as their biofungicide effect against four phytopathogenic fungi.

The application of different extracts produced from different organs of *Pistacia* species have shown bioherbicidal effects. In this line, we firstly studied the direct effect of two extract concentrations (2.5 and 10%) on seed germination and the results indicated that the highest concentrations reduced germination and epicotyl and radicle growth in almost all the weeds and crops studied (*Lactuca serriola, Solanum nigrum, Lolium rigidum, Zea mays, Triticum aestivum*, and *Lens culinaris*). Nevertheless, the 2.5% extract did not cause reduction in the germination of the narrow-leaf weed *Lolium rigidum* and all the studied crops, indicating the selective and dose-dependent effect. According to these results, Bulut et al. [31] showed an inhibitory effect when 200 ppm of *Pistacia terebinthus* extract was applied, while no differences were found at lower concentrations. Furthermore, different studies showed the bioherbicidal effect of *Pistacia khinjuk* and *Pistacia vera* leaf extracts in different weed species such as *Amaranthus retroflexus, Chenopodium album, Physalis alkekengi, Sonchus arvensis, Vicia narbonensis* and *Lolium rigidum* [13-16]. Alyousef and Ibrahim [12] studied the inhibitor effect of *Pistacia vera* hull powder in *Medicago sativa, Diplotaxis erucoides, Sonchus arvensis* and *Papaver hybridum*. In this line, in the present study we further studied the effect of the application of 10% hull extract and hull powder in pots with vermiculite, which corresponds to a dose of 0,025 kg/m^2^ of pistachio hulls and 0,25 l/m^2^ of hulls’ extract. The results showed that no effect was produced in lentil and maize crops, but an inhibition was shown in weeds. *Solanum nigrum* development was reduced under the application of both bioherbicides formats, while *Lolium rigidum* was only inhibited when ground hull was applied. All the studies seem to confirm a more noticeable bioherbicide effect against broad-leaf than narrow-leaf weeds. Curiously, *Lactuca serriola* development was reduced under extract application in lentil assay, while no differences were observed in maize assay. Despite no differences between the treatments, a reduction in *Lactuca serriola* development was observed in the corn assay compared to the lentil assay. This could indicate that corn cultivation itself seems to have an allelopathic effect on this broad-leaf plant [32]. As it was shown there is a selectivity between monocots and dicots weed species, which could be related to plant selection by allelopathy affecting some species more than others [33].

Allelopathy is a natural phenomenon in which an organism affects negatively or positively the functioning of other organisms by releasing secondary metabolites [34]. The allelopathic compounds are mainly secondary metabolites that are secreted into the rhizosphere and affect the development of close plants delaying or inhibiting seed germination and growth [35]. Among these compounds, alcohols, fatty acids, phenolic compounds, flavonoids, terpenoids and steroids can be found. Depending on the botanical compound and weeds, plant allelochemicals have different modes of action based on activities such as repellency, growth suppression, protein denaturation or respiratory impairment [36].

*Pistacia vera* hull constitutes the predominant part of pistachio by-product which contains a high content of these bioactive compounds [21]. These authors pointed out that the phenolic compounds identified in pistachios belong to three groups of phenolic acids, flavonoids, and tannins, which agreed with the results of our extracts. In pistachio hulls, Barreca et al. [11] highlighted the presence of gallic acid, 4-hydroxybenzoic acid, and protocatechuic acid as the most abundant hydroxybenzoic acids. Similar results were described by Erşan et al. [37], highlighting the presence of gallic acid and monogalloyl glucoside in pistachio hulls. Bellocco et al. [38] in pistachio hulls identified the presence of cyanindin-3-O-glucoside and cyanidin-3-O-galactoside. The cyanindin-3-O-glucoside has been pointed out as the major anthocyanin in pistachio hulls [39]. A large variation in the anthocyanins of mature pistachio hulls has been described, however, all of them appear to be cyanindins [37]. Phenolic compounds present different solubility depending on the extraction method [40] which together with cultivar and production area could explain the differences found in different studies. It has been previously demonstrated that water is the most efficient solvent to obtain the higher concentration of phenolic compounds from pistachio leaf and shell organs [41]. In accordance with our results, gallic acid and protocatechuic acid had already been reported in many studied related to pistachio hull polyphenols [11, 37, 42]. Several authors agree in pointing out gallic acid as the predominant phenolic compound in pistachio hulls, being found both free and as part of hydrolysable tannins [21, 43]. Thus, in the present study, the most predominant phenolic compounds (gallic and protocatechuic acids) were tested *in vitro*, showing that the application of both pure compounds in an independent manner (using concentration found in pistachio hull 10% extract) notably reduced epicotyl and radicle lengths of the studied weeds. Protocatechuid acid achieved the highest bioherbicidal effect. Similar results were obtained by Reigosa et al. [44] and Guevara-González et al. [45] who showed an inhibition of germination by the application of gallic acid in aqueous solution in different crops. Li et al. [46] investigated the effect of a *Delonix regia* extract, which contains high concentrations of protocatechuic and gallic acid (among other phenolic compounds), on *Lactuca sativa* and *Brassica chinensis* showing a clear inhibiting effect on plant growth of both species. In addition, in our study, weed seeds exposed to both phenolic compounds displayed an oxidised appearance. This effect was also observed by Malerva-Díaz et al. [47] who reported phenolic oxidation when *Metopium brownei* leaf extracts were applied to *Raphanus sativus*. It should be noted that the aqueous leaf extract of *P. histerophorus* rich in phenolic acids avoid embryo growing or caused its death due to problems in mitosis process, suggesting a possible action mechanism of these compounds [48].

Due to the high content of secondary metabolites with antimicrobial effect [14, 21, 41), pistachio hulls could be considered as biofungicide. Biofungicides are increasingly recognized as a sustainable and economically viable alternative for use in agriculture, avoiding the problems caused by chemically synthesized fungicides. However, their effect depends on the plant from which they are obtained and the concentration [49]. In the present study, no *in vitro* effect on the pathogens *Alternaria alternata, Botryosphaeria dothidea, Aspergillus niger* and *Rhizoctonia solani* was observed despite the fact the study was carried out using a high concentration of pistachio hulls (extract 25%). Nevertheless, several authors have observed a suppressive effect on other pathogens. For example, Contreras [50] observed that the application of methanolic extracts of pistachio (*Pistacia lentiscus*) and rose (*Cowania plicata*) exhibited fungicidal activity on *Colletotrichum coccodes* and *Fusarium oxysporum*, respectively. In addition, several phenolic compounds such as gallic or protocatequic acids can also inhibit the growth of different pathogens. El-Nagar et al. [51] reported that the application of gallic acid and two of its derivatives produced fungistatic action and inhibited the mycelial radial growth of *Alternaria solani* in tomato plants in a dose-dependent manner. In addition, it has been demonstrated that gallic acid derivatives also play an antifungal activity against *Fusarium graminearum* in wheat plants [52]. And protocatequic derivatives exhibited in vitro and in vivo antifungal activity against *B. cinerea* disturbing the integrity of fungi cell membrane and causing oxidative stress and membrane lipid peroxidation [53]. Altogether indicated that pistachio hull extract has a selective fungicidal effect.

## 5. Conclusions

Pistachio hulls are considered agricultural by-products that have been shown to have an allelopathic effect that makes them potential candidates for a role as bioherbicides in sustainable agriculture. In this study, the application of extract and powder coming from pistachio hulls have demonstrated allelopathic effects against *Solanum nigrum, Lactuca serriola* and *Lolium rigidum* weeds in *in vitro* and *in vivo* assays being the most important bioherbicide effect against broad-leaf weeds. Powder seems to be a more efficient strategy to inhibit *Lolium rigidum* weed. The allelopathic effects are mainly due to the high concentration of phenolic compounds such as gallic and protocatequic acids, which were found in the highest concentration, since the pure compounds application inhibits seed germination and development. Nevertheless, no biofungicide effect has been revealed against the studied pathogenic fungi. More studies are required in soil and field conditions to elucidate the mode of action of these biochemicals in order to find more effective and greener strategies to combat weeds and pathogens.

## Acknowledgments

This research was funded by the Grant (2-2022) “DESAFIO Universidad-Empresa” Program, Junta de Castilla y León, co-funded by the European Regional Development Fund (ERDF) and the Junta de Castilla y León.

